# Chromosome Number of the Monarch Butterfly, *Danaus plexippus* (Linnaeus 1758) and the Danainae

**DOI:** 10.1101/107144

**Authors:** Christopher A. Hamm

**Author notes:** Current address: College of Veterinary Medicine, University of California, Davis, One Shields Avenue, Davis, California 95616.

## Abstract

The monarch butterfly, *Danaus plexippus* (Linnaeus 1758) is a charismatic butterfly known for its multi-generational migration in Eastern North America, and is emerging as an important system in the study of evolutionary genetics. Assigning genes or genetic elements to chromosomes is an important step when considering how genomes evolved. The monarch butterfly has a reported haploid chromosome number of 30. The specimens that this count is based on are from Madras, India where *D. plexippus* does not occur. As a consequence the reported haplotype for the *D. plexippus* is incorrect. A literature review revealed that the specimens this count was based on were most likely *Danaus genutia* (Cramer 1779), which Linnaeus and others conflated with *D. plexippus*. In a 1954 Opinion (#282) the International Code of Zoological Nomenclature decided that the name *D. genutia* refers to specimens originating from the Orient and *D. plexippus* to those from North America. To clarify the haplotype number of *D. plexippus* from North America I conducted chromosome squashes of male 5th instar larvae. All specimens examined had a haploid chromosome count of n = 28. Following these new data, I then reconstructed the evolution of haplotypes throughout the Danainae subfamily to provide a testable hypothesis on the evolution of chromosome number for this group.

## Introduction

The monarch butterfly, *Danaus plexippus* (Linnaeus 1758) (Lepidoptera: Nymphalidae) is a large, charismatic insect native to North America and famous for its multi-generational migration across the eastern part of its range (Fig 1.). The monarch is emerging as an important system in the study of evolutionary genetics, and has had a draft of its genome published (Zhan et al. 2011) that has led to significant insights into the evolution of migration and aposematic coloration in butterflies (Zhan et al. 2014). Ultimately, assigning genes to chromosomes will allow for important insights into the evolution of chromosome number evolution and synteny (conserved ordering of genes on homologous chromosomes).

**Figure 1.**
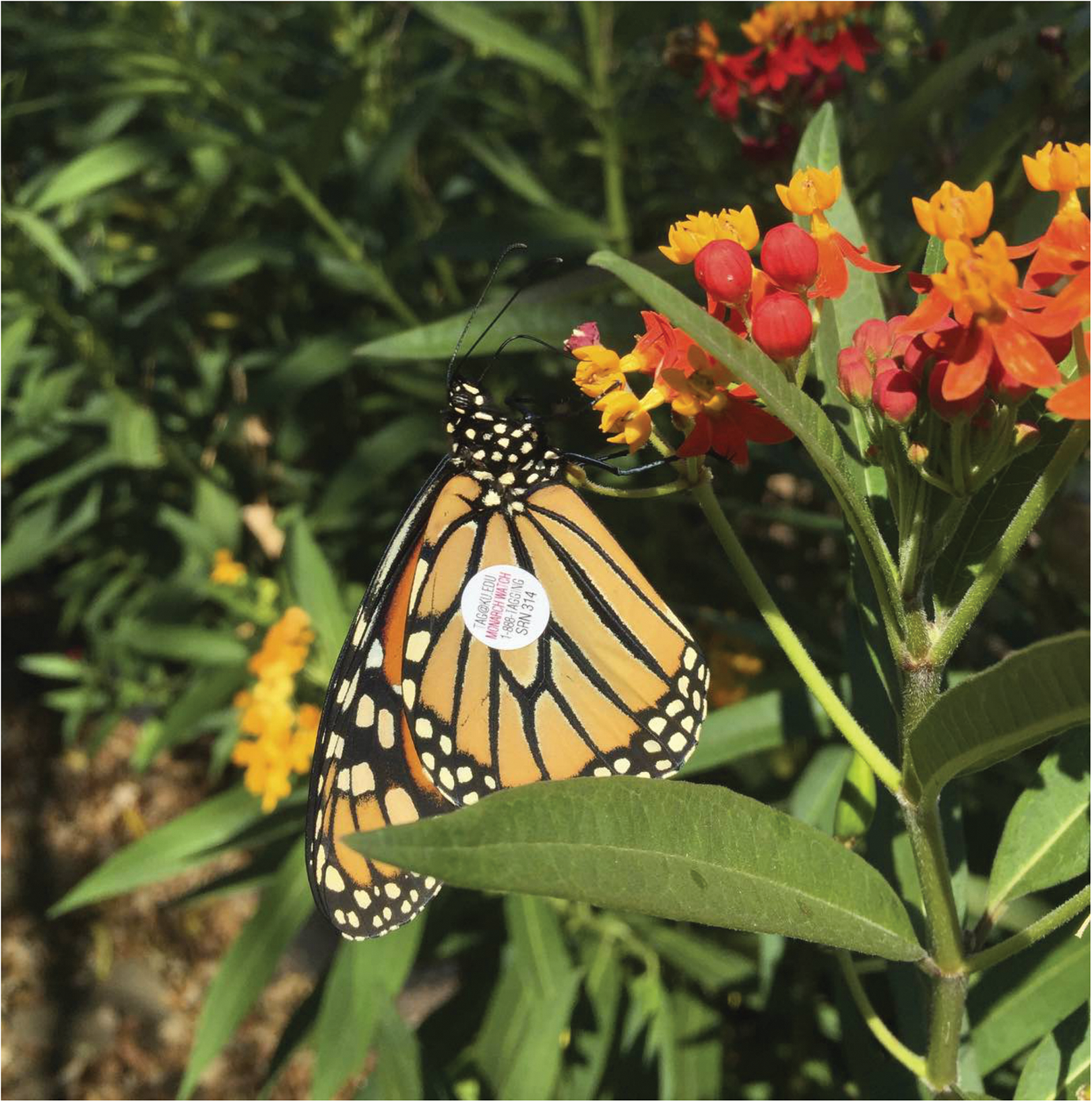
Photo of Monarch Watch tagged female *Danaus plexippus* nectaring on *Asclepias curassavica* in Kansas.

Researchers investigating the evolution of chromosome number, particularly those working in the Lepidoptera, are keen to have accurate counts so that ancestral states can be reconstructed with some accuracy. This is especially important in the Lepidoptera, which have holokinetic chromosomes that allow the attachment of spindle fibers at multiple locations along the length of each body (White 1973). Multiple attachment points are thought to allow the chromosomes of Lepidoptera break into many small fragments (Robinson 1971; White 1973). As such, there can be significant variation in chromosome number within a Family, Genus or species. For example *Eucheira socialis* Westwood 1834 (Lepidoptera: Pieridae) exhibits a haploid chromosome count range from n = 24 to n = 39 (Underwood et al. 2005), and *Eunica malvina* Bates 1864 (Lepidoptera: Nymphalidae) has a range of n = 14 o n = 31 (Brown et al. 2007).While there are clearly species that exhibit a range of haploid numbers, it appears there is strong selection to maintain a particular haploid number of chromosomes that vary by taxa (White 1978; Ahola et al. 2014).

Researchers investigating genomics of *D. plexippus* and reporting its karyotype have cited Brown et al. 2004, which identifies the haploid chromosome count as n = 30. Reading Table 4 from Brown et al. 2004 reveals the specimens that generated this number were from Madras, India and the karyotype number was generated by Rao and Murty 1975. There are no *D. plexippus* native to India. It is likely that Rao and Murty 1975 did not karyotype the North American monarch butterfly *D. plexippus*, but rather *Danaus genutia* (Cramer 1779) (Lepidoptera: Nymphalidae). There has been significant confusion over the proper name of the monarch-like butterfly that occurs on the Indian subcontinent, to the extent that Linnaeus conflated the species. The confusion was removed by Opinion 282 of the International Code of Zoological Nomenclature, which established the name *D. genutia* shall refer to specimens originating from the Orient and *D. plexippus* shall refer to specimens from North America (ICZN Opinion 282) (ICZN 1954; Brown 1968). Thus, there has been no karyotyping of *D. plexippus.* To fill the gap left by this revelation, I conducted chromosome squashes of male *D. plexippus* reared from wild caught butterflies by Monarch Watch at the University of Kanas.

## Materials and Methods

To establish karyotypes for *D. plexippus* from Kansas I prepared chromosome squashes following the protocol used by Underwood et al. 2005. Briefly, I removed the testes from 5th instar larvae and placed them into fixative. The testes were then macerated, spread over a glass microscope slide, dried and stained with Giemsa. A total of two larvae were initially screened for chromosome counts, and one week later 4 additional specimens were examined for a total of 6. I then used a Zeiss Axio Imager 2 compound microscope with Zeiss EC Plan-Neofluor 100x oil immersion lens to identify cells that were arrested at meiotic metaphase I or II. I used an attached AxioCam MRc5 CCD camera and captured images with ZEN 2012 (Blue Edition) Imaging suite. Because intra-individual variation in haplotype count is know (Pearse and Ehrlich 1979) each specimen was scored a minimum of four times (four technical replicates from the same individual).

Using these new data, I reconstructed haplotype evolution in a phylogenetic context. Phylogenies that contain taxa in the Danaini have been published but did not suit my purposes because they were either constructed at the genus level (Wahlberg *et al.* 2009) or were constructed using morphological characters (Brower *et al.* 2010). I downloaded the sequences referenced in Brower et al. (2010) (one mtDNA locus, two nDNA loci) and the corresponding sequences of *Libythea celtis* to serve as an outgroup. The sequences were aligned by locus using Geneious v6.1.8 (Kearse et al. 2012) and fit to models of molecular evolution with MrModelTest v2.3 (Nylander 2004) run in PAUP* v4.1b10 (Swofford 2001). I then used the program MrBayes v3.2.5 (Ronquist and Huelsenbeck 2003) to construct a phylogeny where each locus was allowed a variable evolutionary rate that selected by Akaike Information Criterion (Akaike 1974) in MrModeltest. The Metrolpolis-Hastings Markov Chain Monte Carlo (MCMC) analysis was run with 10 chains (9 heated, one cold) for 1 million generations and the chains were sampled every 5,000 generations. Convergence of the chains was visually assessed by comparing log likelihood values and ensuring that the standard deviation of split frequencies was below 0.01. I removed 25% of the remaining 500 trees as burn-in and calculated posterior probabilities based on the post-burn-in trees.

Data for haplotypes count in the Danaini and *L. celtis* were acquired from synthetic literature (Robinson 1971; Brown et al. 2004) and confirmed with relevant the primary literature. Using the phylogenetic trees created previously, I pruned the tree using functions in the R package geiger (Pennell et al. 2014) so that only tips with character data remained. Chromosome counts in the Lepidoptera feature high levels of dysploidy, (variation in chromosome number by a single value that takes two forms), ascending dysploidy (addition of a single chromosome due to fission) and descending dysploidy (loss of a single chromosome through fusion). As such, chromosome counts cannot be treated as continuous variables and require alternative approaches (Mayrose et al. 2010). The pruned tree and data were used to model chromosome evolution with the program `chromEvol` (Glick and Mayrose 2014), which employs a maximum-likelihood approach to evaluate models of chromosome evolution across a phylogenetic tree. I used the available chromosome count data to provide `chromEvol` with count frequencies when dysploidy was present in species, and fit a variety of models to the data and phylogeny. Specifically, I considered three models: 1) constant rates of ascending (λ) and descending (δ) dysploidy; 2) linear rates of ascending (λ_l_) and descending (δ_l_) dysploidy where rates are conditional on the current chromosome number; and 3) a model with all four parameters (λ, δ, λ_l_, δ_l_). I did not consider any models that allowed demi-ploidy or genome duplications, as I am aware of no evidence indicating that these phenomena have been observed in the Lepidoptera (White 1973). I fit these models to the data and phylogeny using a range of starting parameters (1.0 to 10^-7^) to minimize the possibility of local optima, and used 10 optimization steps followed by 10,000 simulations to compute the expected number of changes by transition type and compared models using AIC (Akaike 1974). All data and code have been archived on GitHub (https://github.com/butterflyology/Monarch_haplotype) with Zenodo (https://zenodo.org/badge/latestdoi/81117728) and the microscopy images have been archived with FigShare (https://figshare.com/articles/Monarch_images/4633390).

## Results and Discussion

I observed a haplotype count for every *D. plexippus* male was n = 28 (Figure 2), which is two below the count reported for what we now know to be *D. genutia* (Rao and Murty 1975). These chromosomes appear similar in shape to other danaines previously reported (Figure 2) (Brown et al. 2004) and this haplotype count is still consistent with other counts within *Danaus* (Table 1).

**Figure 2.**
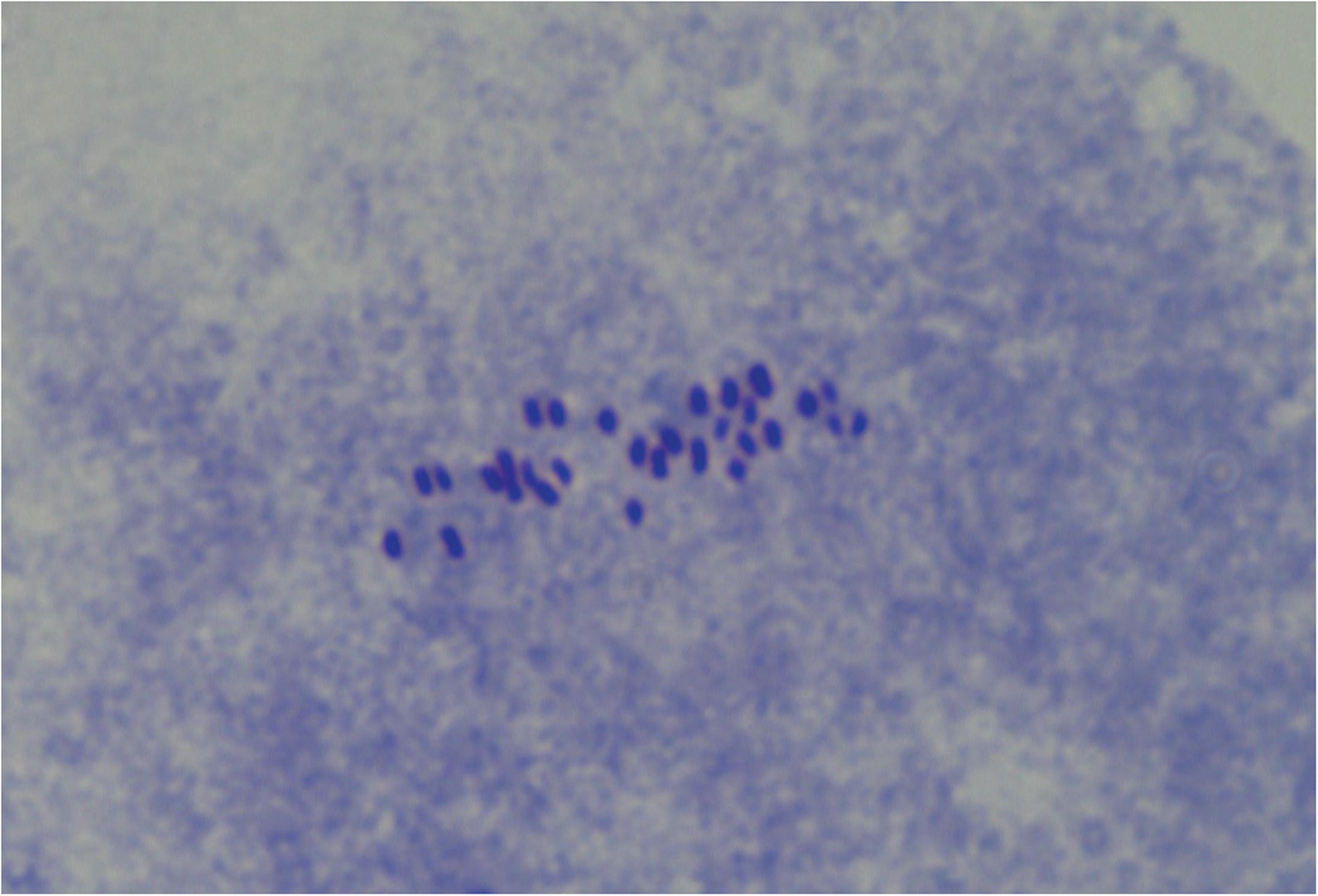
Photo of chromosome squash from male *Danaus plexippus* taken at 1000X magnification through oil-immersion lens on a XYZ microscope.

**Table 1.** Chromosome counts for *Danaus* species and appropriate reference.

| *Danaus* | Haploid count | Reference |
| --- | --- | --- |
| *cleophile* | 30 | Brown et al. 2004 |
| *chrysippus* | 30-31 | Srivastava and Gupta 1961; de Lesse and<br>Condamin 1962 |
| *eresimus* | 30 | Brown et al. 2004 |
| *erippus* | 30 | Brown et al. 2004 |
| *genutis* | 30 | Rao and Murty 1975 |
| *gilippus* | 29 | Brown et al 2004 |
| *limniace* | 41-46 | Bernardi and de Lesse 1964 |
| *plexaure* | 30-31 | Brown et al. 2004 |
| *plexippus* | 28 | This study |

I modeled ploidy changes in the Danaini tribe in a phylogenetic context. Lepidopteran chromosomes are holokinetic, i.e. kinetochores are located along the length of the chromosome, rather than concentrated at one location, which is thought to allow spindle fibers to attach at multiple locations along a chromosome and therefore increase the rate of fission (White 1973), resulting in multiple chromosome numbers within genera, species, and even individuals (Robinson 1971; Underwood et al. 2005; Dincă et al. 2011; Šíchová et al. 2015). This feature, known as disploidy, occurs throughout the Lepidoptera and is present in the Danaini (Maeki 1961; Brown et al. 2004).

I reconstructed a fully resolved and extremely supported phylogenetic tree of the Danainini that had the same topology as Wahlberg et al. (2009) and Brower et al. (2010). Each of the three chromEvol models we examined converged to similar parameter estimates regardless of starting values, suggesting the algorithm was not stuck on a local optimum. All models generated parameter estimates that predicted both gain and loss of chromosomes in the Danaini, indicating that chromosome fission and fusions have occurred throughout the lineage (Table 2), and reconstructed nearly identical ancestral chromosome counts for nodes, with the only difference the being the estimated count between *L. celtis* and *Tellervo zoilus* (Figure 3). Importantly, a chromosome count of N = 31 is inferred for all ancestral nodes leading to the *Danaus* genus, within which a fusion event leads to an N = 30 count. Subsequent fusions reduce *D. plexippus* to N = 28 and some *D. gilippus* and *D. eresimus* to N = 29.

**Table 2.** chromEvol parameter estimates by model with AIC scores.

| Model | $\lambda$ | $\delta$ | $\lambda_l$ | $\delta_l$ | AIC |
| --- | --- | --- | --- | --- | --- |
| constant | 65.5 | 50.7 |  |  | 104.8 |
| linear |  |  | 2.0 | 2.0 | 105.8 |
| constant + linear | 34.1 | 85.3 | 0.9 | 0.7 | 108.6 |

**Figure 3.**
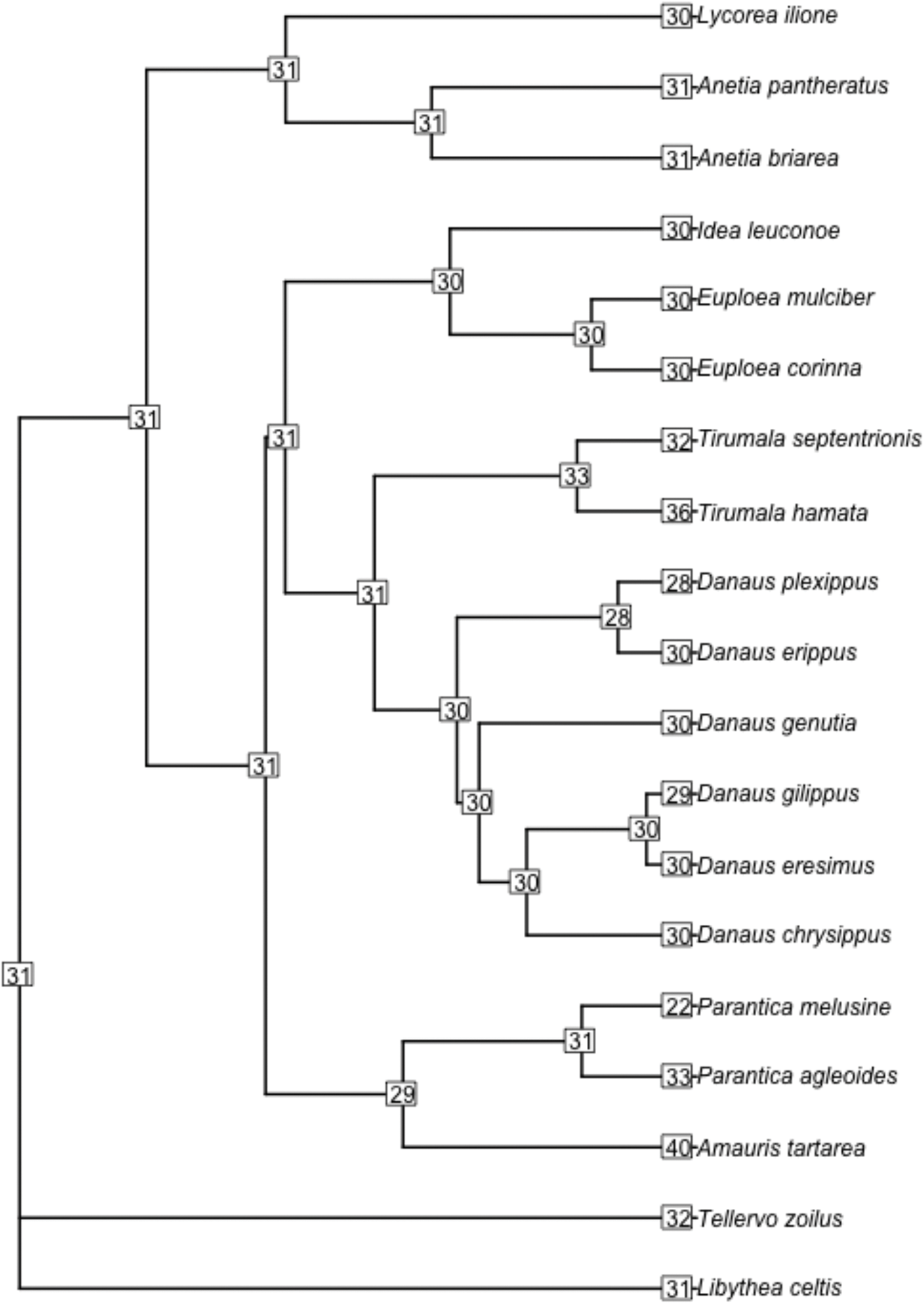
Phylogeny of the Danainae and `chromEvol` reconstruction of haplotype evolution.

This revised chromosome count suggests that minor modification to the results of (Ahola et al. 2014) and their reconstruction of chromosome number evolution in the Papilionoidea. Because chromosome number can vary geographically I strongly encourage the investigation of karyotype in other populations of *D. plexippus* and the Danainae.

## Acknowledgements

I wish to thank Dr. Dessie Underwood (California State University, Long Beach) for teaching me the laboratory techniques used to identify chromosomes, and Dr. Chip Taylor (Monarch Watch, University of Kansas) for providing me with larvae from their rearing project. I also wish to thank Dr. Ben Sikes (University of Kansas) for allowing me use of his lab’s microscope, and Az Klymiuk (University of Kansas) for training me in the use of the microscope. I also wish to thank Dr. Krushnamegh Kunte (National Center for Biological Sciences) for confirming my suspicion about the synonymy.

